# The 5-methylcytosine DNA glycosylase ROS1 antagonizes parent-of-origin specific DNA methylation in Arabidopsis endosperm

**DOI:** 10.1101/2024.11.05.622036

**Authors:** Elizabeth A. Hemenway, Mary Gehring

## Abstract

DNA methylation patterning is a consequence of opposing activities of DNA methyltransferases and DNA demethylases. In flowering plants, two distinct female gametes, the egg cell and the central cell, are fertilized, producing what will become the embryo and the endosperm of the seed. In Arabidopsis, a 5-methylcytosine DNA glycosylase, DME, demethylates regions in the central cell genome, leading to methylation differences between maternally- and paternally-inherited endosperm genomes after fertilization. *DME* is required for endosperm gene imprinting. Homologues of *DME* include *ROS1*, *DML2* and *DML3*. It is unknown whether any of these DNA glycosylases are required for endosperm methylation patterning. We show that *ROS1* prevents hypermethylation of paternally-inherited alleles in the endosperm at regions that lack maternal or paternal-allele methylation in wild-type. Thus, *ROS1* promotes epigenetic symmetry between genomes in the endosperm by preventing paternal genome hypermethylation. We propose that *ROS1* and *DME* act in a parent-of-origin-specific manner at shared endosperm targets, and consider implications for the evolution of imprinting mechanisms.

## Introduction

The presence of the modified base 5-methylcytosine (5mC), DNA methylation, is a common feature of eukaryotic genomes. DNA methylation is an epigenetic mark that can be stably inherited through cell divisions and across generations. DNA methylation represses potentially harmful genomic elements such as repeats and transposable elements (TEs), and can also contribute to transcriptional regulation of protein coding-genes. Pathways that add and maintain 5mC and pathways that remove 5mC each contribute to genomic DNA methylation patterning. In plants, cytosines are methylated in the CG, CHG, and CHH sequence contexts, where H is an A, T or C. A family of bifunctional DNA glycosylases act as DNA demethylases by cleaving the glycosidic bond between the 5mC base and the sugar-phosphate backbone, initiating base-excision repair (BER)^1–3^. Member genes in *Arabidopsis thaliana* are *DME*, *ROS1*, *DML2*, and *DML3*. Homologues of the *DME* family have been found to have analogous function in other flowering plant species^4–7^. At common target regions, *ROS1* opposes the RNA-directed DNA methylation pathway (RdDM), which establishes *de novo* DNA methylation in all sequence contexts and maintains a portion of asymmetric CHH methylation^8–11^. Other pathways also maintain DNA methylation. MET1, a homolog of mammalian DNMT1, is recruited to hemi-methylated DNA, and maintains methylation in the CG sequence context by methylating the unmethylated strand^12–14^. CHG and some CHH methylation is maintained through positive feedback between the maintenance DNA methyltransferases CMT3/CMT2 and H3K9me2, a histone modification established by SET-domain H3K9 methyltransferases SUVH4/5/6^15–19^

DNA demethylase targets have been identified in various plant tissues and life stages by whole-genome DNA methylation profiling in plants carrying mutations in DNA demethylases. *ROS1* preferentially removes DNA methylation from the 5’ and 3’ ends of genes. Many of these demethylated regions correspond TE ends that are near genes, and targeted by RdDM^10^. *DML2* and *DML3* are more lowly expressed in vegetative tissues and are largely redundant with *ROS1* in, but do have unique target sites^9,20^. *DME* has also been shown to contribute to DNA demethylation in vegetative tissues^21^, but primarily has been studied for its major function during the reproductive phase of the plant life cycle, as *DME* is essential for seed viability^22^. *DME* is active in the central cell and is required for maternal allele demethylation of a subset of TEs, typically short fragments, in the endosperm^3,6,23–25^. *DME*/*ROS1* homologs have also been found to be important for seed development in rice and in maize^4,6,7,26^. While the role of *DME* in reproduction has been appreciated for some time, the function of *ROS1*, *DML2*, and *DML3* in plant reproductive tissues has not been extensively investigated. *ROS1* and *DME* have been shown to promote male fertility in Arabidopsis^27,28^. *ROS1* and *DME*-mediated DNA demethylation of pollen-specific genes in the vegetative nucleus is required for their expression and promotes proper pollen tube function^28^.

The endosperm, a critical seed tissue that supports embryo development, has a unique genetic and epigenetic landscape. Resulting from fertilization of the diploid central cell, triploid nuclei of the endosperm contain a 2:1 maternal to paternal genome ratio. Endosperm DNA is hypomethylated relative to that in other tissues, in part due to *DME* expression in the central cell prior to fertilization, as well as repression of methylation pathways^23–25,29^.

The endosperm is the site of gene imprinting, which refers to biased gene expression from either the maternal alleles (maternally expressed imprinted genes – MEGs) or the paternal allele (paternally expressed imprinted genes – PEGs) after fertilization. *DME*-mediated DNA demethylation in the central cell results in maternally-inherited endosperm genomes that are hypomethylated relative to the paternally-inherited endosperm genome, which is necessary for gene imprinting^3,24,30^. However, biased expression of imprinted genes is not always clearly associated with observed differential methylation between maternal and paternal genomes in the endosperm, and various epigenetic states have been identified for imprinted genes^31^. MEGs tend to have high levels of H3K27me3 on the paternal allele, deposited by the PRC2 complex. PEGs are more often associated with paternal genome hypermethylation in their 5’ and 3’ regions, and can be associated with H3K27me3, H3K9me2, and CHG methylation in the gene body of the maternal allele^32–34^.

In Arabidopsis, there is some evidence for *ROS1* activity in seeds. *ROS1* promotes the expression of the imprinted gene *DOGL4* in the endosperm and embryo^35^. *ROS1* is expressed in dry seeds, and regions dependent on *ROS1* for wild-type methylation levels have been identified in whole dry seeds^36^. *DME*, however, is the only demethylase essential for endosperm development in *Arabidopsis thaliana*. In this study, we investigate the role of *ROS1* and *RDD*, if any, in global DNA methylation patterning of the endosperm. We find that *ROS1* contributes to DNA methylation patterning in the endosperm, predominantly by preventing hypermethylation of the paternal genome. We identify regions in the endosperm that are likely co-targeted by *DME* and *ROS1*, where *ROS1* promotes a biallelically-demethylated state in the endosperm.

## Results

### *ROS1* contributes to endosperm DNA methylation patterning

Gene expression datasets indicate that all four 5-methylcytosine DNA glycosylases are expressed in endosperm^37^ (Supplemental Fig. 1A). To determine whether 5-methylcytosine DNA glycosylases other than *DME* contribute to the endosperm DNA methylation landscape, we profiled DNA methylation in the absence of *ROS1* and in the absence *ROS1*, *DML2* and *DML3*. We performed enzymatic-methyl sequencing (EM-seq) on three replicates each of wild-type Col-0, *ros1*, and *ros1-3 dml2-1 dml3-1* endosperm. For *ros1* profiling, we used *ros1-3* mutants, which have a T-DNA insertion in exon 7, and *ros1-*7 mutants, which carry a missense allele in which an essential glutamic acid residue in the DNA glycosylase catalytic domain is changed to a lysine residue^9,38^. We used two different mutant alleles of *ros1* so that we could confirm any observations were not line-specific. Based on previous studies, we predicted that *ros1*-7 is a hypomorphic allele, whereas *ros1*-3 is a null allele. For all samples, endosperm nuclei were isolated from whole seeds at 7 days after pollination (DAP) by FANS based on their triploid DNA content (Supplemental Fig. 1B). We obtained methylomes of 98 to 194x genome coverage (Supplemental Table 1). To facilitate direct comparison between endosperm and a vegetative tissue, we also profiled methylation in three replicates of wild-type Col-0, *ros1-*3, and *ros1-7* rosette leaves, obtaining 2C and 4C nuclei by FANS. Globally, the total fraction of methylated cytosines was not significantly different between *ros1-3*, *ros1-7*, nor *rdd* and wild-type in the endosperm (Fig. 1A, p-value ≥ 0.02). This result is consistent with previous findings in *ros1* vegetative tissues^9^. To identify potential discrete regions of differential methylation we used Dispersion Shrinkage for Sequencing Data (DSS) to identify differentially methylated regions (DMRs) between the demethylase mutant endosperm and wild-type in the CG, CHG, or CHH methylation sequence contexts^39,40^ (Fig. 1B,C). The *ros1-3* mutation is linked to regions of the Ws ecotype genome remaining on chromosome 2; these regions were removed from analysis in *ros1-3*, *ros1-7*, and Col-0 prior to calling DMRs to avoid over-calling false-positive DMRs^9,21^. For identifying DMRs between *rdd* and Col-0, we removed regions of the Ws genome remaining on chromosomes 2 and 3, linked to *ros1-3* and *dml2,* respectively^9,21^. Using this method, we identified 1,624 total DMRs in any sequence context between *ros1-3* and Col-0, 913 total DMRs between *ros1-7* and Col-0, and 1,319 total DMRs between *rdd* and Col-0 (Supplemental Table 2). The DMRs were short, with the most abundant fraction being 50-100 bp in length (Supplemental Fig. 2A). By genome browsing, short DMRs called by DSS are often surrounded by regions that also appear differentially-methylated, but are not identified as a DMR. Thus, we are likely underestimating the true degree of differential methylation among genotypes (Fig. 1B).

**Figure 1:**
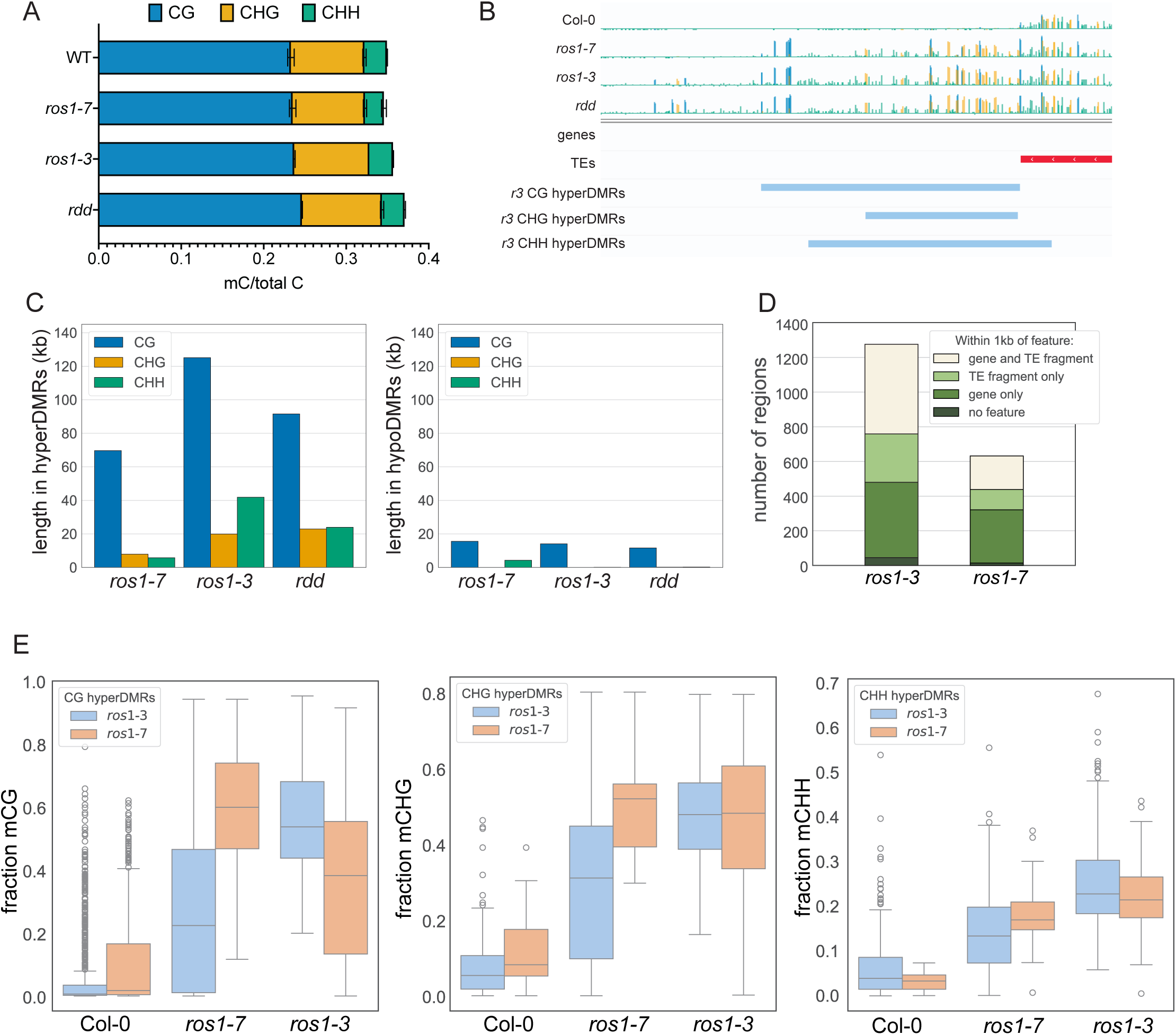
*ROS1* prevents endosperm hypermethylation. A) Fraction of cytosines by sequence context that are methylated in Col-0, *ros1-7*, *ros1-3*, and *rdd* in 3C and 6C endosperm nuclei. Error bars represent standard deviation from the mean. B) Example of a region (around Chr5:9,271,400-9,272,200) with observable DNA hypermethylation in all demethylase mutant backgrounds in the endosperm. C)Total length of hyperDMRs (left) and hypoDMRs (right) by sequence context in each mutant. D) The number of ROS1 endosperm target regions (after merging hyperDMRs identified across sequence contexts into a single list of hypermethylated regions) found within 1kb of or intersecting a feature. E) Methylation of *ros1-3* (blue) or *ros1-7* (orange) hypermethylated regions in WT, *ros1-7* or *ros1-3* endosperm. Weighted average mC for each *ros1* hyperDMR by sequence context. One biological replicate of each genotype is presented, additional replicates presented in Supplemental Figure 3.

Consistent with the molecular function of DNA demethylases, most DMRs were more highly methylated in demethylase mutant endosperm compared to wild-type endosperm (Fig. 1C). To assess the replicability of *ROS1* target regions across the unique mutant allele genotypes, we calculated the level of DNA methylation at hyper-DMRs defined using either *ros1-7* or *ros1-3* in all sequenced genotypes (Fig. 1E, Supplemental Fig. 3). Across replicates and samples, *ros1* mutants are more highly methylated than Col-0 at DMRs identified in any *ros1* mutant background. These results support a high degree of similarity between the two *ros1* mutant gentoypes. The fewer DMRs identified between *ros1-7* and Col-0 compared to *ros1-3* and Col-0, support *ros1-7* being a hypomorphic allele whereas *ros1-3* is a null allele. Finally, these results show that disruption of *DML2* and *DML3* cause only minor increases in endosperm hypermethylation compared to loss of *ROS1* alone (Fig. 1C, Supplemental Fig. 3). Thus, we focused our additional analyses and experiments on *ROS1*.

### *ROS1* prevents DNA methylation spreading from target TEs in the endosperm

*ROS1* is known to maintain DNA methylation boundaries at target TEs, preventing aberrant DNA methylation ‘spread’^10,11,41^. The relative hypo-methylation of the wild-type endosperm genome relative to leaf and seedling tissue prompted us to further investigate the impact of *ROS1* at TE boundaries. We merged *ros1-3 vs*. Col-0 hyperDMRs called in different sequence contexts into a single list of *ROS1* target regions to identify broad features of *ROS1* endosperm targets. The same merging was performed for *ros1-7 vs.* Col-0 hyperDMRs. Consistent with previous results, *ROS1* target regions in the endosperm are often found near TEs; over half of *ROS1* target regions identified in the *ros1-3* mutant endosperm were within 1 kb or intersecting a TE (Fig. 1D). We identified 1463 out of the 34856 Araport11-annotated TE fragments as within 1 kb of or intersecting a *ROS1* target region defined by *ros1-3* methylation data, and 624 TE fragments within 1 kb of or intersecting a *ROS1* target region defined by *ros1-7* methylation data. For subsequent analysis, we merged directly adjacent or overlapping TE fragments, resulting in 19536 genomic TE ‘regions’, 1051 of which were associated with a *ros1-3* hypermethylated region. We calculated average percent mC in 100 bp windows around TE ends for *ROS1* target-proximal TEs and for all non-*ROS1* target-proximal TEs (Fig. 2A, Supplemental Fig. 4-5). In Col-0 endosperm, TEs that are near a *ROS1* target region are less methylated in flanking regions (average 26% CG methylation) than TEs not proximal to a *ROS1* target (average 38% methylation), with a sharper boundary between methylation in the flanking region and in the body of the TE. We observed DNA methylation spreading up to approximately 1 kb outside the annotated boundary of *ROS1* target-associated TE regions in *ros1* mutant endosperm. We conclude the loss of *ROS1* degrades sharp TE methylation boundaries in the endosperm.

**Figure 2:**
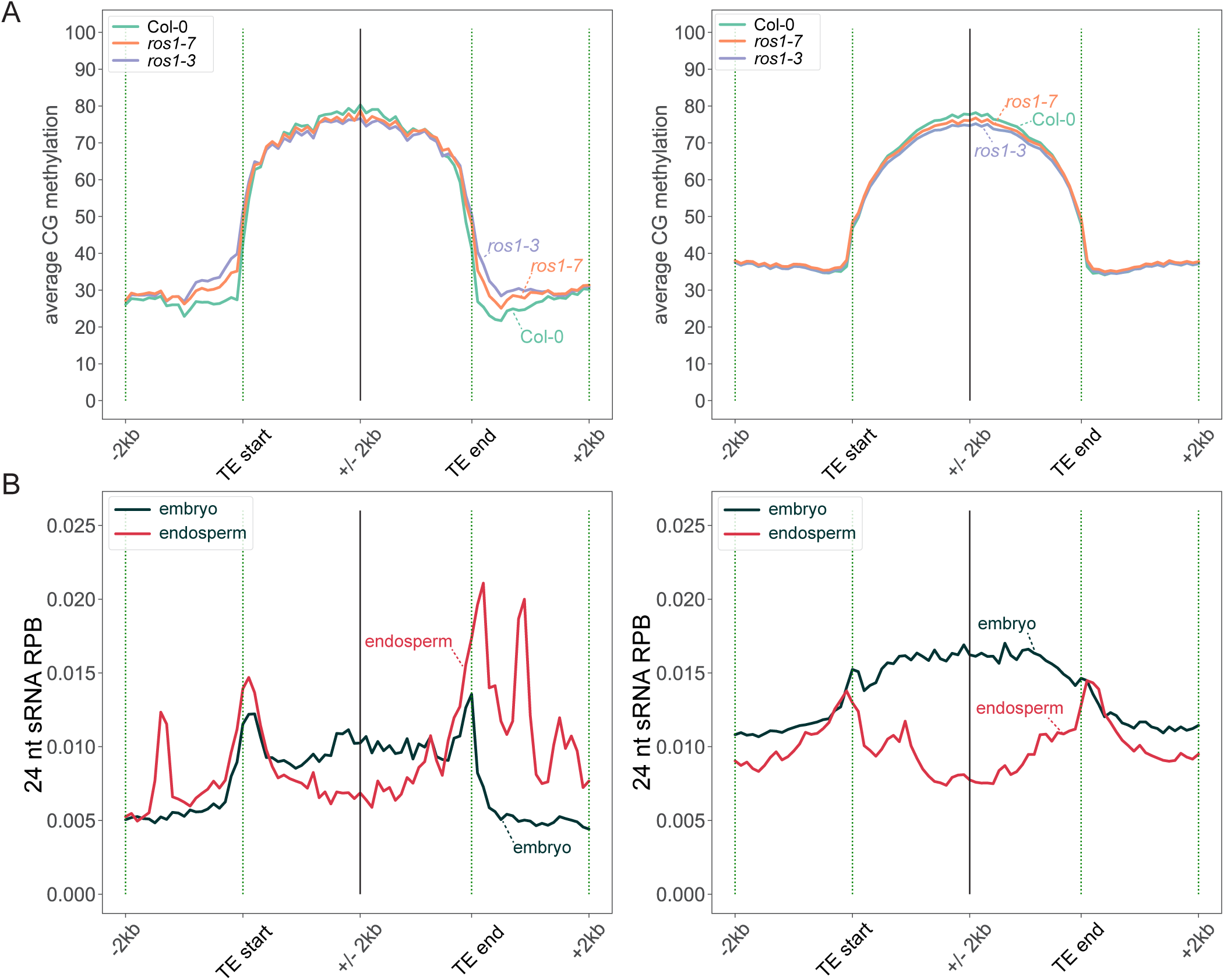
Flanking regions of *ROS1* target-associated TEs are hypomethylated but enriched for 24 nt sRNAs in wild-type endosperm. A) Average percent CG methylation determined in 100 bp windows 2kb outside and 2kb inside of TE regions. TEs that are within 1kb of a *ROS1* endosperm hyperDMR are defined as *ROS1* target-associated TEs (left; n=1169). TEs not within 1kb of a *ROS1* endosperm-defined target region are on the right (n=18367). B) Average coverage (plotted as reads per base (RPB)) of wild-type embryo and endosperm 24 nt sRNAs for each in 50bp windows 2kb outside and 2kb inside of TE regions.

We examined whether regions of methylation spreading were associated with 24-nucleotide small RNAs, which participate in RdDM, in wild-type Col-0^42^. The bodies of *ROS1* target-proximal endosperm TEs produced fewer 24 nt sRNAs in wild-type embryos than non-*ROS1* targeted TEs^42^ (Fig. 2B, black lines). This finding is consistent with *ROS1* acting at TEs in euchromatic regions, as it has been shown that TEs which produce more sRNAs in the endosperm relative to the embryo are depleted from pericentromeric heterochromatin^10,11,41,42^. The proximal regions of *ROS1* target-proximal TEs, where methylation spreads in *ros1* endosperm, produced more 24-nt sRNAs in wild-type Col-0 endosperm than in the embryo (Fig. 2B). These findings suggest that in wild-type, *ROS1* counteracts methylation directed by tissue-specific small RNA production at TE ends in the endosperm.

### *ROS1* targets have reduced capacity for hypermethylation in endosperm

To determine whether *ROS1* acts at unique sites in the endosperm, we investigated the extent to which DNA hypermethylation at *ros1* vs Col-0 hyperDMRs was tissue-specific. We calculated the level of cytosine methylation in our Col-0 and *ros1-3* leaf methylation data as well as in published data from wild-type Col-0 and *ros1* sperm cells^28^ for regions defined as endosperm *ros1-3* hyperDMRs. Sperm DNA methylation was of particular interest because the endosperm is a product of fertilization between a sperm and the central cell. We found that regions defined as endosperm *ros1-3* hyperDMRs were also hypermethylated in *ros1-3* mutant leaf and sperm relative to the respective wild-type tissue (Fig. 3A, Supplemental Figures S3 and S6). In wild-type plant tissues, *ros1-3* hyperDMRs reflect the same DNA methylation features that have been observed at a genome-wide scale: wild-type endosperm has lower DNA methylation levels than does leaf, and sperm has higher DNA methylation levels relative to leaf or endosperm (Fig. 3A, Supplemental Figure 6). TEs are known to be hypomethylated in the endosperm relative to non-endosperm tissue^23,24^, and we investigated whether this might represent an endosperm-specific role of *ROS1* at target TEs. However, we found that, in their bodies, *ROS1*-target TEs were indeed hypomethylated in the endosperm relative to leaf tissue, but that this TE-body hypomethylation was not *ROS1*-dependent (Fig. 3B).

**Figure 3:**
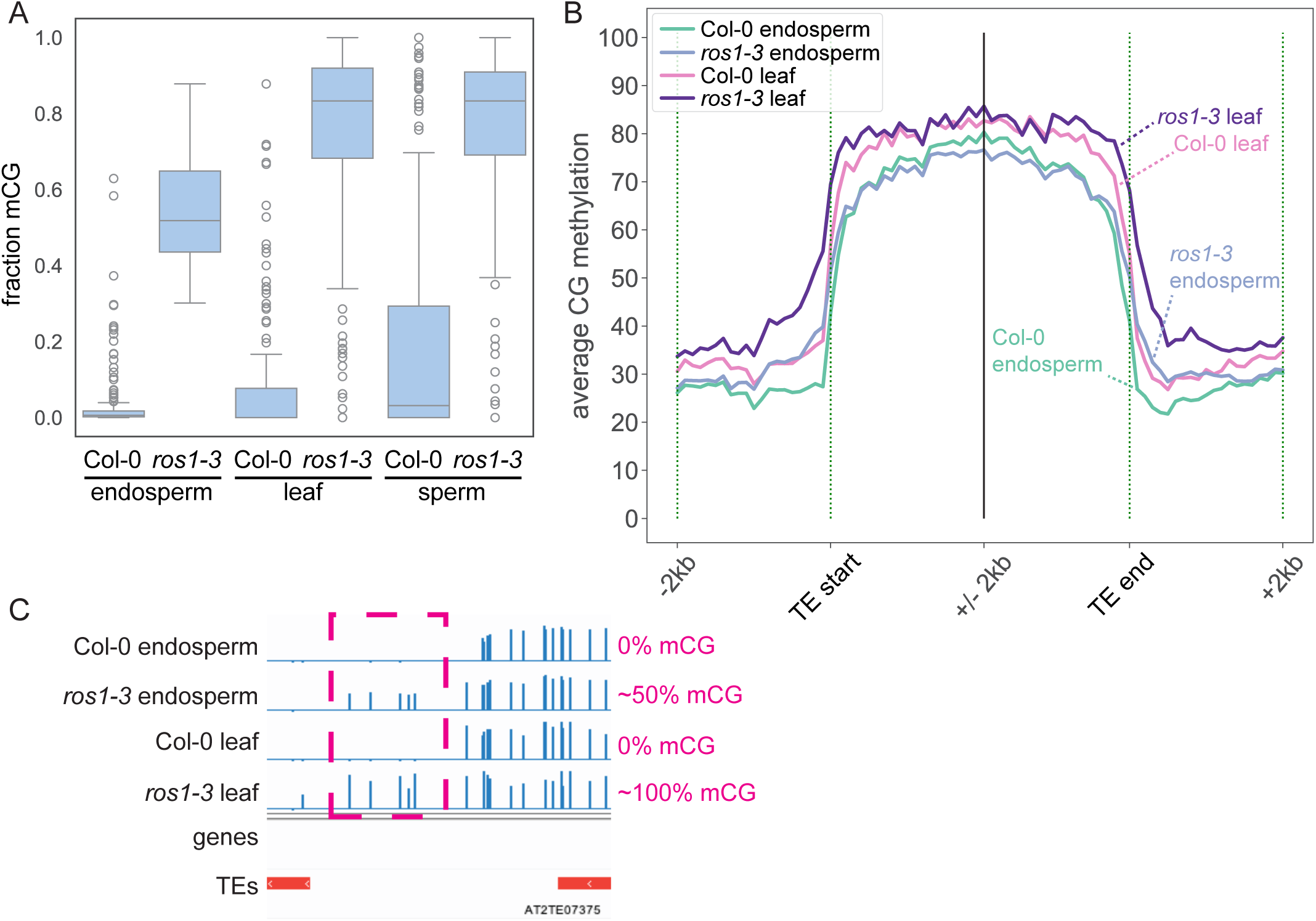
*ROS1* target regions display limited hypermethylation in endosperm relative to mutant leaf or sperm. A) Weighted average mCG in *ros1-3* CG hyperDMRs defined in the endosperm. All wild-types are Col-0. Sperm data from Khouider et al 2021. B) Average percent mCG in 100 bp windows 2 kb inside and outside of merged TE regions within 1 kb or intersecting a *ROS1* target region defined in *ros1-3* vs Col-0, in any sequence context. C) Genome browsing example of a *ROS1* target region with limited hypermethylation in *ros1-3* mutant endosperm relative to *ros1-3* mutant leaf.

Although *ROS1* endosperm targets are also *ROS1* targets in leaf and sperm, they display overall lower levels of CG DNA methylation in both Col-0 and in the *ros1* mutant (Fig. 3A,B). Additionally, we observed a subset of *ros1* endosperm CG-hyper DMRs with a substantially reduced capacity for hypermethylation in endosperm relative to leaves. By genome browsing, these regions were characterized by low or no CG methylation in both wild-type leaf and endosperm, close to 100% methylation in *ros1* mutant leaf, but closer to 50% methylation in *ros1* mutant endosperm (Fig. 3C). To identify the more extreme examples of this phenomenon, we filtered *ros1* endosperm CG-hyper DMRs for regions that were 50% as methylated in *ros1* mutant endosperm as in *ros1* mutant leaf; 15.75% of *ros1-3* endosperm CG hyperDMRs met these criteria and we refer to these regions as hypermethylation-limited CG DMRs. This DMR type was also observed, but to a lesser extent, in *ros1-7*, where 6.14% of endosperm CG hyper-DMRs met this threshold (Supplemental Table 3).

What underlies the failure to reach the fully hypermethylated state in *ros1* endosperm? We considered two possible non-mutually-exclusive possibilities: 1) these regions are variably methylated between nuclei of the *ros1* endosperm or 2) these regions are differently methylated between the maternal and paternal genomes in the *ros1* endosperm. In support of the second hypothesis, *ROS1* has been show to prevent hypermethylation of the paternal allele of the *DOGL4* promoter in endosperm^35^, and in our dataset CG sites in the *DOGL4* promoter are consistently less methylated in the *ros1* mutant endosperm relative to leaf (Supplemental Figure 7).

### *ROS1* precludes hypermethylation on the endosperm paternal genome at target loci

To test if *ROS1* targets are differentially methylated between paternal and maternal genomes we performed allele-specific whole-genome EM-seq using F1 endosperm isolated from reciprocal crosses between *ros1* mutants in the Col-0 (*ros1-3*) and C24 (*ros1-1*)^1^ backgrounds, along with appropriate controls. Endosperm nuclei were collected from three replicates each of Col-0 x C24, C24 x Col-0, *ros1-3* x *ros1-1*, *ros1-1* x *ros1-3*, C24 x C24 and *ros1-1* x *ros1-1* (female parent in cross written first). SNPs between C24 and Col-0 were used to assign reads to a parent-of-origin after sequencing^43^ (Supplemental Table 1).

We compared the methylation of maternal and paternal alleles at the previously-identified *ROS1* target regions in wild-type and *ros1* endosperm. In the wild type, CG methylation of maternal and paternal alleles at *ROS1*-target regions is highly correlated (Pearson’s *r* = 0.83) (Fig. 4A). In contrast, CG methylation of maternal and paternal alleles of the same regions in *ros1* endosperm is not correlated (Pearson’s r=-0.06) (Fig. 4B). The lack of correlation is caused by the specific gain of paternal allele methylation in *ros1* endosperm (Fig. 4A-C). DMRs that we previously defined as especially hypermethylation-limited are characterized by particularly low CG methylation on the maternal allele in both wild-type and *ros1* mutant endosperm, approximately 20% mCG on the maternal allele in the *ros1* mutant (Fig. 4A-B, orange dots). The phenomenon of paternal allele hypermethylation is replicable across ecotypes, as we observed a comparable paternal bias on the C24 genomes at CG DMRs called between the *ros1-1* and wild-type C24 endosperm (Supplemental Fig. 8). Non-CG hyperDMRs are more biallelically hypermethylated than CG hyperDMRs although a reduced correlation between maternal and paternal alleles is also observed in comparisons of Col-0 to *ros1-3* endosperm (Supplemental Fig. 9). The observed paternal bias of *ROS1* activity is variable between sequence contexts.

**Figure 4:**
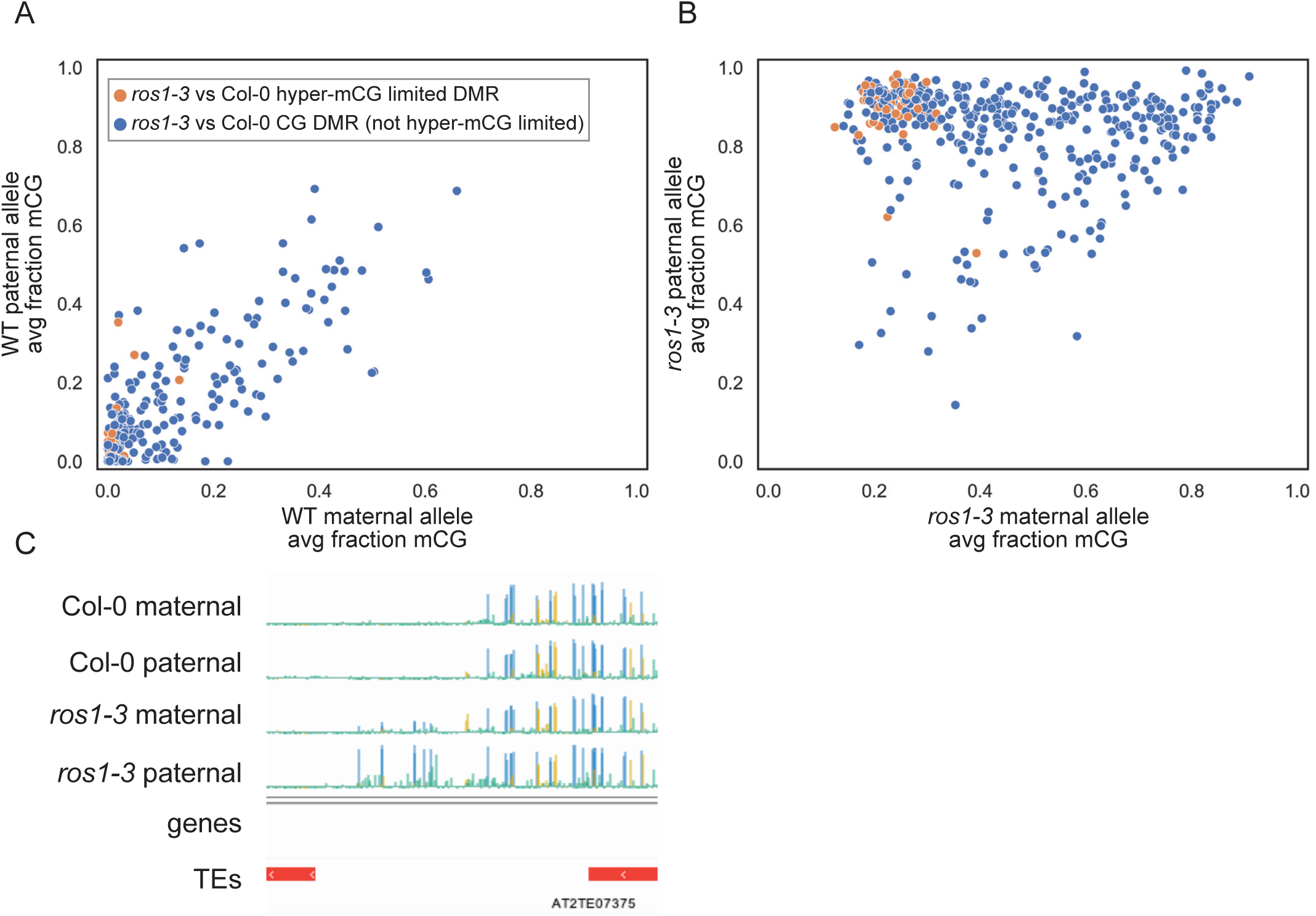
Hypermethylation in *ros1* mutant endosperm is biased for the paternal allele. A) Weighted averaged fraction mCG levels for *ros1-3* CG hyperDMRs on maternal and paternal genomes in F1 endosperm isolated from Col x C24 (WT maternal) C24 x Col (WT paternal), B) *ros1-3* x *ros1-1* (*ros1-3* maternal), and *ros1-1* x *ros1-3* (*ros1-3* paternal). Values plotted are the averaged weighted sum value across biological replicates. Orange dots represent hypermethylation-limited CG DMRs identified in the non-allelic data, which display particularly low levels of maternal methylation relative to paternal methylation in *ros1-3*. C) Genome browser example of the *ros1* target region displayed in Figure 3C, now with distinguished maternal and paternal alleles.

Studies of ROS1 activity *in vitro* have not conclusively revealed a sequence context preference of ROS1 demethylase activity, and *in vivo* whole-genome sequencing data has repeatedly shown that *ROS1* prevents hypermethylation of cytosines in all sequence contexts^2,9,44^. Considering these results, we propose the differences observed between sequence contexts indicate *ROS1* target regions are differentially targeted by methylation-establishing and maintenance pathways on the maternal and paternal genomes in the endosperm. Overall, *ROS1* prevents hypermethylation of paternal allele more strongly than the maternal allele in the endosperm, particularly in the CG sequence context.

### *ROS1* promotes a bi-allelically demethylated state

To gain a more complete sense of the potential relationships between paternally-acting *ROS1* pre- and post-fertilization, we also identified DMRs between maternal and paternal genomes within a wild-type background and within a *ros1* mutant background (Fig. 5A). For each genotype, DMRs were called in mCG, mCHG, and mCHH sequence contexts independently, but resulting maternally-hypomethylated DMRs across sequence contexts were merged together into one list of regions per genotype for subsequent analyses. Previously, it was shown that endosperm maternal allele hypomethylation depends, at least in part, on the activity of maternal *DME* before fertilization^3,6,23–25^. Regions (n=1586) that are demethylated on the maternal allele and methylated on the paternal allele in F1 progeny endosperm of all four reciprocal crosses (WT x WT, *ros1* x r*os1*) are likely regions where *DME* establishes a parental DNA methylation difference by acting in the central cell pre-fertilization, and this difference is maintained independently of *ROS1* (Fig. 5A, Supplemental Table 4). Regions defined as maternally-hypomethylated in both *ros1* x *ros1* cross directions, but neither WT x WT cross direction were also identified (Fig. 5A,B, Supplemental Table 4). These regions (n=274) are biallelically-demethylated in the wild-type endosperm, and gain DNA methylation relative to wild-type predominantly on the paternal allele in the absence of *ROS1*. The hypomethylation of the maternal allele relative to the paternal allele is observable in the *ros1* mutant background at these regions. This suggests that in wild-type, the maternal allele lacks methylation due to a *ROS1*-independent mechanism, whereas the paternal allele is demethylated by *ROS1* (Fig. 5A,B). The presence of such regions indicates a role of *ROS1* in preventing differential DNA methylation between maternal and paternal genomes in the endosperm, specifically differential methylation where the paternal allele is more highly methylated than the maternal allele.

**Figure 5:**
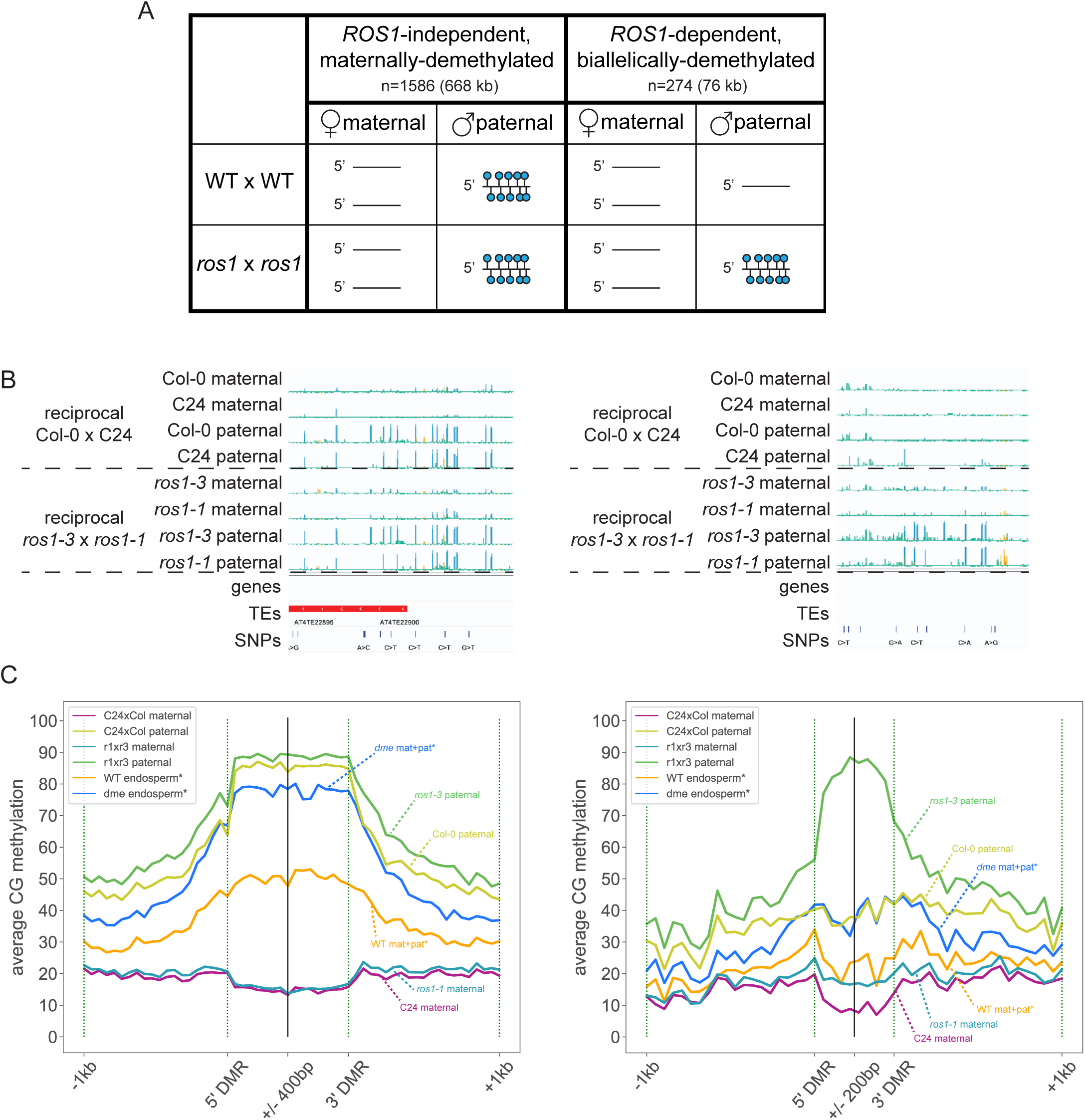
*ROS1* promotes biallelic demethylation by preventing hypermethylation of the paternal allele at some targets. A) A cartoon diagram depicting simplified examples of presented maternally-hypomethylated DMR classes as we have defined them, with corresponding quantification of these regions from our analyses. B) Genome browser examples of the presented maternally-hypomethylated DMR classes. C) CG methylation of maternal and paternal genomes from the present study from selected biological replicates (all replicates plotted in Supplemental Fig.10) across 50bp windows, 400 bp inside and 1 kb outside each end of maternally-demethylated, *ROS1*-independent regions (left) and 200 bp inside and 1 kb outside each end of *ROS1*-dependent regions (right). We used different numbers of windows inside the two classes of DMRs to account for the differences in average length between them (Supplemental Fig. 2).

We hypothesize regions that are maternally-demethylated in wild-type endosperm, and are not dependent on *ROS1* for proper demethylation are targeted by *DME* in the central cell prior to fertilization. To test this hypothesis, we examined methylation levels in these regions using published non-allelic *dme* endosperm data^25^. We observed hypermethylation of our proposed *DME*-targeted, maternally-demethylated regions in *dme-2* endosperm data that represents combined maternal and paternal genome methylation. We also examined whether maternal allele hypomethylation at *ROS1*-dependent, biallelically-demethylated regions is dependent on *DME*. We observed hypermethylation in *dme* endosperm at *ROS1*-dependent biallelically-demethylated regions (Fig. 5C, Supplemental Fig. 10), but to a lesser extent (3-5% CG methylation gain in *ros1*) than *ROS1*-independent regions. This suggests that *DME* prevents hypermethylation of the maternal allele at *ROS1*-dependent regions, but that other factors may also be involved. For example, it is possible that methyltransferases, such as *MET1*, do not maintain methylation of the maternal allele of these regions in the central cell and early endosperm, which would also prevent maternal allele methylation, even in the absence of *ROS1.* Additionally, we find that maternally-demethylated DMR-proximal regions are more highly methylated in wild-type endosperm than are *ROS1*-dependent, biallelically-demethylated DMR-proximal regions (Fig. 5C), most notably on the paternal allele, and that maternally-demethylated DMRs are longer than biallelically-demethylated DMRs (Supplemental Fig. 2). These observations indicate a difference between *ROS1*-independent, presumed *DME*-only targets and *ROS1*-dependent, presumed *DME* and *ROS1* targets, which could contribute to parent-of-origin specific methylation in wild-type endosperm.

### Inheritance of wild-type *ROS1* does not fully rescue hypermethylation in F1 heterozygous endosperm

Several possible mechanisms could explain the observed biased effect of *ROS1* on endosperm paternal allele methylation state. One is paternal allele-specific demethylation of these regions in the endosperm after fertilization. Another, not mutually-exclusive, possibility is that demethylation by *ROS1* is not allele-specific in the endosperm but that there is no methylation on the maternal allele to be removed. Alternatively, *ROS1* could act prior to fertilization in the male germline, leading to inheritance of a demethylated paternal allele in the endosperm. Using available data of demethylase mutant and wild-type sperm cells as well as our own data from leaf, we observed that *ROS1*-dependent, biallelically-demethylated regions are hypermethylated in *ros1* mutant sperm and leaf to the same degree that they are methylated on the paternal allele in the endosperm in the CG sequence context (Supplemental Fig. 11).

Based on our comparisons between leaf, sperm, and endosperm DNA methylation data, we sought to test if *ROS1* is active at *ros1*-hypermethylated regions after fertilization. More specifically, is paternal allele hypermethylation at *ROS1* targets rescued in the presence of a wild-type *ROS1* allele? We hypothesized that if *ROS1* is acting only through the male germline, then a wild-type paternal *ROS1* allele would be necessary for bi-allelic demethylation at *ROS1*-dependent, biallelically-demethylated regions. To test this hypothesis, we performed allele-specific methylation profiling on endosperm of F1 seed from reciprocal crosses between *ros1-3* (in the Col-0 background) and wild-type C24 (Fig. 6A). In this design, the endosperm is heterozygous for the *ros1* mutation but either the maternal or paternal sporophytic tissues and gametophytes are without *ROS1*. If *ROS1* activity is required before fertilization in the male germline to cause hypomethylation of paternal alleles after fertilization, then regions paternally hypermethylated in homozygous mutant *ros1-3* endosperm (Fig. 5) will also be paternally hypermethylated in heterozygous *ros1-3* endosperm when the mutation is inherited through the paternal parent. In contrast, maternal inheritance of *ros1* should not cause paternal allele hypermethylation in endosperm. If *ROS1* is instead acting after fertilization to demethylate paternal alleles in endosperm, then paternal allele hypermethylation will not be observed in heterozygous *ros1-3* endosperm when the mutation is inherited through the paternal parent.

**Figure 6:**
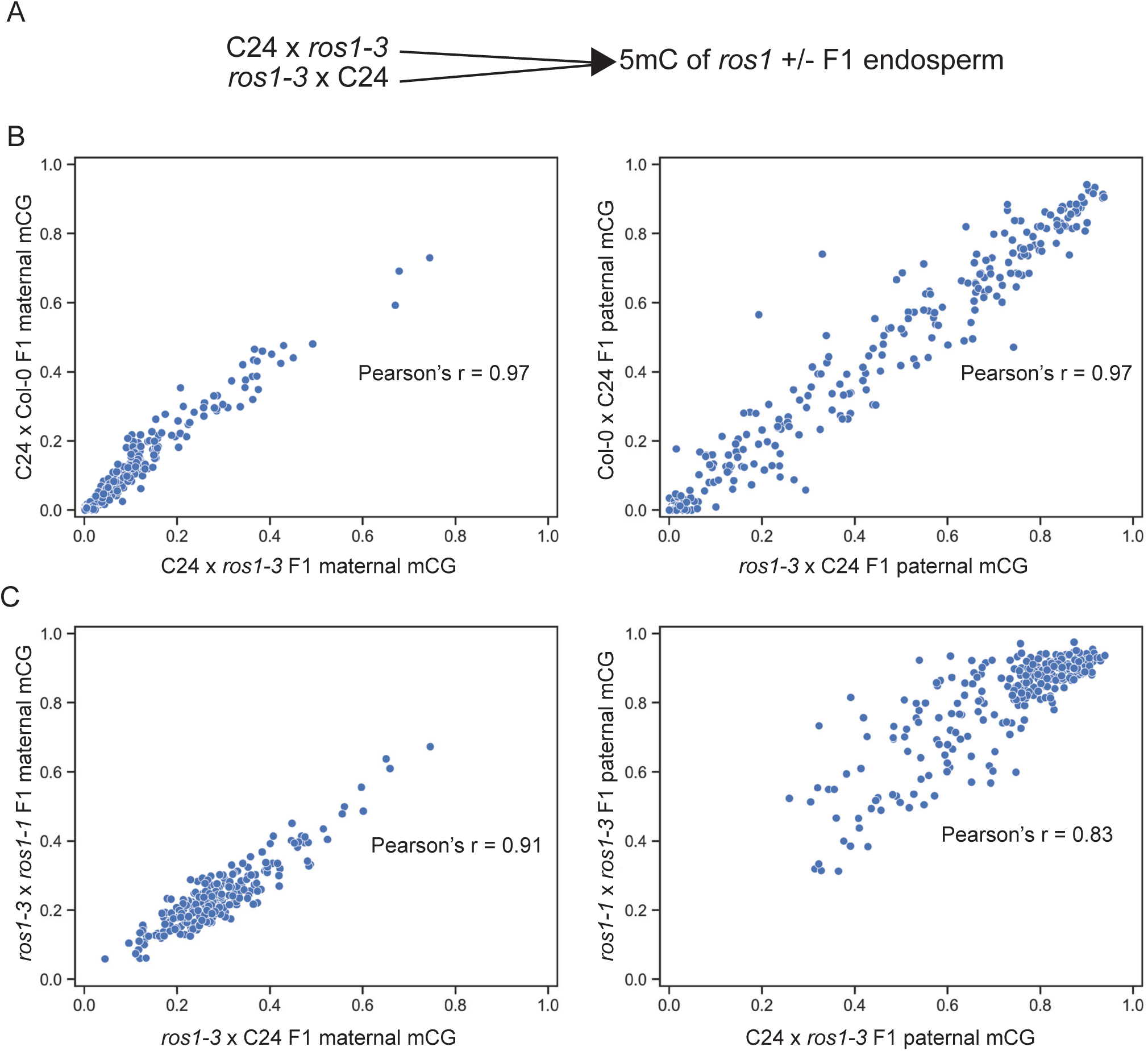
Inheritance of paternal *ros1-3* in the endosperm is sufficient for most, but not complete, paternal allele hypermethylation. A) Graphical depiction of experimental design. To compare genomes inherited from a wild-type *ROS1* background to genomes inherited from a mutant *ros1* background in the endosperm, we reciprocally crossed wild-type C24 and *ros1-3*. Then, we sequenced the endosperm DNA from F1 progeny which are heterozygous for a wild-type copy of *ROS1*. B) Weighted averaged mCG values of identified biallelically-demethylated, *ROS1*-dependent regions identified in the previous experiment. Comparing averaged values across biological replicates calculated from alleles in the wild-type C24 genomes of a *ROS1* homozygous background to those of a heterozygous *ros1-3/ROS1* mutant background. Pearson’s r values for both plots have a p-value of less than 1.0E-100. C) As in B., but now comparing values calculated from alleles in the *ros1-3* genomes of a *ros1* homozygous background to a heterozygous *ros1-3/ROS1* mutant background. Pearson’s r values for both plots have a p-value of less than 1.0E-70.

**Figure 7:**
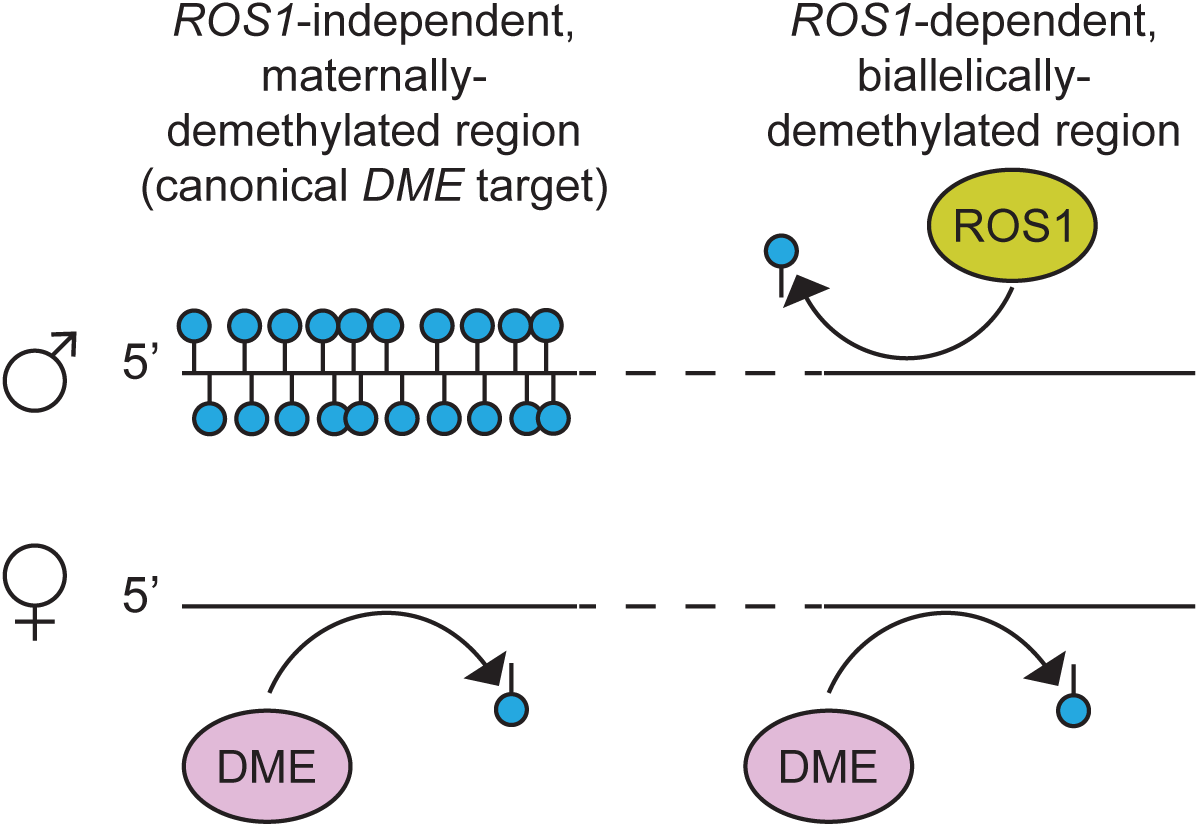
Model for ROS1 activity at maternally-hypomethylated regions in wild-type endosperm. In this model, ROS1 removes DNA methylation from the the paternal allele at *ROS1*-dependent, biallelically-demethylated regions, but not at *ROS1*-independent, maternally-demethylated regions. Most paternal-specific DNA demethylation by ROS1 is likely occuring prior to fertilization. DME is presumably acting to prevent hypermethylation of the maternal allele at *ROS1*-independent regions, and may likely be preventing maternal hypermethylation at *ROS1*-dependent regions.

We compared the CG methylation levels at biallelically-demethylated regions in the F1 homozygous progeny of reciprocal crosses to the respective mutant genome in the C24 by *ros1-3* F1 heterozygotes. There was a high correlation between methylation of C24 maternal alleles in wild-type (C24 x Col-0) and *ros1* heterozygous backgrounds (C24 x *ros1-3* Col-0) (Pearson’s r=0.97) (Fig. 6B). Maternal allele methylation was also highly correlated between homozygous mutant (*ros1-3* x *ros1-1*) and heterozygous (*ros1-3* x C24) endosperm, although to a lesser degree (Pearson’s r = 0.91) (Fig. 6C). These data indicate that maternal allele methylation remains low in these regions regardless of the *ROS1* genotype. The paternal genome in a homozygous *ros1* mutant background (*ros1-3* x *ros1-1*) and the paternal genome in a heterozygous *ros1*/+ background where *ros1* was inherited from the paternal parent (*ros1-3* x C24) had the lowest correlation (Pearson’s r = 0.83) (Fig. 6C). The lower correlation was caused by higher methylation of *ROS1* target regions in *ros1* homozygous mutant endosperm compared to *ros1* heterozygous endosperm (Fig. 6C).

Overall, we conclude that the majority of paternal allele hypermethylation at *ROS1*-dependent, biallelically demethylated regions in *ros1* mutant endosperm is inherited through the male germline. However, we observe a slight reduction in paternal DNA methylation at *ROS1* target regions in the *ros1*/+ F1 heterozygous endosperm (Fig. 6C), indicative of some active maternal *ROS1* in the endosperm after fertilization, which is able to partially rescue paternal CG hypermethylation.

## Discussion

In *Arabidopsis thaliana*, loss of DNA demethylase activity is well tolerated under normal lab growth conditions, except during female gametogenesis that gives rise to endosperm. The endosperm is uniquely sensitive to epigenome disturbances, making it an informative tissue in which to study epigenetic mechanisms. Studies of the DNA demethylase-encoding gene, *DME*, highlight this point; *DME* is required in the female gametophyte for seed development and to establish an asymmetric epigenome state between the maternal and paternal genomes^3,22,45^. Although *ROS1* is not essential for seed development in Arabidopsis, we have characterized a unique role of *ROS1* for maintaining proper epigenetic patterning in the endosperm. We first identified *ROS1*-target region in the endosperm using high-quality, high-coverage whole genome DNA methylomes, generated by enzymatic-methyl sequencing. These regions are also identifiable as *ROS1* targets in leaf data we generated for this study, but are biased for hypermethylation of the paternal allele in a *ros1* mutant endosperm. Thus, a biallelically demethylated state in the endosperm is *ROS1*-dependent at some regions.

We found that loss of *ROS1* leads to a biased gain of DNA methylation on the paternally- inherited genome at *ROS1* target regions in the CG and CHH sequence contexts. Notably, we observe a more biallelic *ROS1-*dependence in the CHG sequence context, which could be attributable to differences in the activity of methylation establishing and/or maintenance pathway activity in the endosperm. We identified regions of maternal hypomethylation in the wild-type ColxC24 reciprocal cross F1 progeny, and based on the known role of *DME* in demethylating the maternal genomes of the endosperm prior to fertilization, as well as our analyses of previously published *dme* data^25^, we hypothesize that these regions are dependent on *DME* for maternal allele demethylation. Methylation status at these regions is largely *ROS1*-independant, although we have observed that there is often some *ROS1*-dependancy at the ends of regions exhibiting maternal hypomethylation (Fig. 5C). We identified an additional, novel class of regions that are biallelically-demethylated in wild-type endosperm, but gain methylation in a *ros1* mutant, primarily on the paternal allele. We propose these regions are dependent on *DME* for maternal allele demethylation, and we show *ROS1* prevents paternal allele hypermethylation at these regions (Fig. 8). Low maternal allele methylation at these regions could also be a result of lack of methyltransferase targeting, which will be an interesting direction for future studies. We observed slight rescue of paternal allele hypermethylation in *ros1* heterozygous endosperm when *ros1* was inherited from the paternal parent (Fig. 6C), suggesting *ROS1* is capable of demethylating the paternal allele in the endosperm after fertilization. However, the majority of paternal hypermethylation of *ROS1*-dependent, biallelically demethylated regions in *ros1* endosperm is inherited through the male germline. One caveat to these results is that *ROS1* expression has been identified as paternally-biased in other studies^32^ (Supplemental Fig. 12), and thus there might be very little ROS1 in the endosperm from a *ros1* mutant paternal parent.

We note that in the *ros1* mutant endosperm, *ROS1*-dependent, biallelically-demethylated regions gain resemblance to regions associated with imprinted genes in wild-type endosperm, with a methylated paternal allele and a maternal allele with low methylation (Fig. 5A). In light of this observation, one intriguing hypothesis is that for a region to gain imprinted status on an evolutionary timescale, it might need to both gain/retain *DME* target status in the female gametophyte but lose *ROS1* target status in the male germline. Future work will investigate the molecular and genetic relationship of *ROS1* and *DME* in reproductive tissues to further our understanding of gene imprinting mechanisms in the seed.

## Methods

### Plant material

Plants for non-allele-specific methylation profiling in endosperm and leaf (Figures 1-3) were grown in a Conviron growth chamber at 22°C using 22 hr light/2hr dark cycles with 120 uM fluorescent light. Plants for allele-specific methylation profiling (Figures 4-6) were grown using 16 hr light/8hr dark cycles; set-point for temperature was 22°C. The mutants *ros1-3* and *ros1-3; dml2-1; dml3-1* were originally described in Penterman et al., 2007^9^. The *ros1-7* allele was described in Williams et al 2015^38^. In the C24 background, the *ros1-1* mutant allele was described in Gong et al 2002 and was obtained from ABRC (CS66099)^1^. Genotypes for all plants were PCR-verified.

### Preparing endosperm samples for EM-seq

Flowers were emasculated and pollinations were preformed 2 days after emasculation. Seven days after pollination (DAP), seeds were removed from 3-6 siliques and placed into 120 μl of Nuclei Extraction Buffer, available as part of the CyStain UV Precise P kit (Partec North America #055002) on ice. Seeds were ground manually using a sterile pestle. 800 μl of companion Nuclei Staining Buffer was added to ground seeds, and mixture was filtered 1-2 times through sterile Cell Trics 30 μm filters and kept on ice. 3C and 6C nuclei were then sorted into 200 μl of Buffer ATL from Qiagen QIAamp DNA micro kit (#56304) with BD FACSAria SORP equipped with an 100 mW UV laser using a nozzle size = 100 microns, Sheath pressure = 20 psi. 20 μl of Proteinase K solution was added to the nuclei mixture and incubated overnight at 56°C. The next day, remaining steps for QIAamp DNA micro kit were performed, following the protocol for “Isolation of Genomic DNA from Tissues”. DNA was eluted in 20 μl of nuclease-free water and kept at −20°C until all samples were prepared.

### Preparing leaf samples for EM-seq

Rosette leaf tissue from 4-week-old plants was collected and chopped manually in Nuclei Extraction Buffer using a razor blade. All following nuclei preparation steps were the same as for seeds. For leaf tissue, 2C and 4C nuclei were collected using the BD FACSAria conditions described above. DNA extraction steps were the same as for endosperm nuclei.

### EM-seq Library Prep

To prepare converted libraries for whole-genome sequencing of DNA methylation, we used the EM-seq kit from NEB (#E7120S). 1 μl diluted 1:100 pUC19 control DNA per 10-30 ng sample DNA was spiked in as per kit instructions. Using a Covaris sonicator, DNA samples for the non-allelic seed and leaf were sheared in 130 μl of 1x TE buffer, 6x16 mm tubes using conditions: [Peak power = 175 W, Duty factor = 10% Cycles/burst = 200 Time = 180 sec]. TE buffer was then exchanged for 50 μl nuclease-free water using a 1X Ampure bead clean up before continuing the EM-seq protocol. Conditions were later optimized, and for the allelic endosperm experiments DNA was sheared in 50 μl of 1x TE buffer, 6x16 mm tubes under conditions [Peak power = 175W Duty factor = 10% Cycles/burst = 200 Time = 80 sec]. All remaining library prep steps were then performed as written in the EM-seq manual; we used 7-8 cycles of PCR amplification to amplify libraries. Libraries were pooled at approximately equimolar concentration using library quantity quantified from Fragment Analyzer readouts prior to a final bead clean-up of remaining adapter fragments, and sequenced on an Illumina NovaSeq S4 machine, 120 x 120 paired end.

### Processing non-allele specific EM-seq data

Data processing was performed based on the Bismark pipeline (https://felixkrueger.github.io/Bismark/)^46^. Briefly, reads were trimmed and filtered using Trim galore using standard settings for phred33 quality control and paired-end Illumina reads (https://github.com/FelixKreger/TrimGalore). Trimmed reads were then mapped to TAIR10 genome containing the pUC19 methylated control sequence using Bismark alignment. The number of mismatches allowed in a seed alignment (--N) was set to 1 and the maximum insert size (--X) to 600. Bismark deduplicate was used to remove PCR duplicates using default settings. Methylation information was extracted from deduplicated reads with Bismark Methylation Extractor using default settings for paired-end reads. Initial conversion and protection quality checks, and analysis of whole-genome methylation status were performed (Supplemental table 1). Where applicable, we excluded the region of annotated Ws homology, Chr2: 8,802,496-15,397,296 from *ros1-3* endosperm and leaf samples and the matched wild-types prior to DMR calling^21^. We removed the same region of Chr 2 as well as the annotated Ws region at Chr3: 688,340-5,117,803 from *rdd* bulk endosperm and matched wild-types prior to DMR calling^21^. Presence or absence of Ws regions did not have an out-sized effect on DMR identification, and we did not remove Ws regions from *ros1-3* data in the allelic analysis for DMR calling.

### C24 pseudogenome generation

To facilitate mapping reads from C24 ecotype genomes in F1 hybrids, we generated a TAIR10 genome with substituted SNPs for the C24 variant^43^. We then prepared this genome for mapping using Bismark genome preparation with default settings.

### Processing allele-specific EM-seq data

Reads were trimmed and filtered using Trim galore using standard settings for phred33 quality control and paired-end Illumina reads. Trimmed reads were then mapped to TAIR10 genome containing the pUC19 methylated control sequence using Bismark alignment. The number of mismatches allowed in a seed alignment (--N) was set to 1 and the maximum insert size (--X) to 600. Bismark deduplicate was used to remove PCR duplicates using default settings. Methylation information was extracted from deduplicated reads using Bismark Methylation Extractor using default settings for paired-end reads. Initial conversion and protection quality checks, and analysis of whole-genome methylation status were performed at this stage. Remaining reads that did not map to TAIR10-pUC19 were then mapped to a pseudo-C24 genome and deduplicated. Mapped reads from both mapping runs were assigned to a parent-of-origin using the SNPs identified between Col-0 and C24 by Jiao et al 2020, accounting for C-T or G-A SNPs, using assign_to_allele.py^47^. Methylation information was then extracted from each parental genome individually using Bismark Methylation Extractor, before combining outputs from the same parental genomes from the two mapping runs.

### Identifying differentially methylated regions (DMRs)

DMRs were called using Dispersion Shrinkage for Sequencing data^39,40^. A cytosine was required to have 3 supporting reads to be included in the DSS analysis. The following parameters were utilized: [DMR.delta=0.3(CG), 0.2(CHG), 0.1(CHH); DMR.p.threshold = 1e-02; DMR.minlen = 100; DMR.minCG = 5 ; DMR.distance.merge = 100 ; DMR.pct.sig = 0.1].

### Merging DMRs

When applicable, we compiled DMRs for each mutant genotype identified in the CG, CHG, and CHH sequence contexts. We used Bedtools merge to merge any overlapping DMRs (which would occur if a region was called as a DMR in more than one sequence context) into one target region.

### Summing methylation across regions of interest

For summed mC analyses, as in figures 1E, 3A, 4A-B, 6B, input methylation data for cytosines with at least 5 informative reads are intersected with regions of interest using Bedtools intersect. Then, a weighted mean methylation level is calculated for each region, weighted by sequencing depth at each cytosine^32^.

### Analysis of average methylation patterns for features of interest

For each feature, average percent mC value is calculated over set windows inside and outside the feature. Window lengths and length surrounding feature ends are described for each analysis in the results. An average is then calculated for each window across the meta- feature and plotted^47^. When performing this analysis using 24 nt sRNA data, an average value of coverage per base pair was used as input data^42^.

## Data availability

Sequencing data is available on GEO (GSE280598). Scripts used to process and analyze data are available on Github (https://github.com/Gehring-Lab/ROS1_endo). Published data sets available through GEO GSE15922, GSE94792, GSE14114 were used and referenced accordingly.

## Acknowledgments

We thank the Whitehead Genome Technology Core and the Whitehead Flow Cytometry Core for experimental support. We thank George Bell for supporting and writing code for the initial implementation of the DSS pipeline. We thank Souraya Khouider, Roman Podolec and Caroline Martin for providing feedback on the manuscript, as well as other members of the Gehring lab for valuable scientific discussion and feedback. Research reported in this publication was supported by the National Institute of General Medical Sciences of the National Institutes of Health under Award Number R35GM145321 to MG. The content is solely the responsibility of the authors and does not necessarily represent the official views of the National Institutes of Health. MG is an Investigator of the Howard Hughes Medical Institute.

## Supplemental Information

Supplemental Table 1: Mapping statistics and conversion rates.

Supplemental Table 2: DMRs identified in endosperm, *ros1-3 vs*. Col-0, *ros1-7 vs*. Col-0, and *rdd vs*. Col-0.

Supplemental Table 3: Hypermethylation-limited mCG DMRs identified in *ros1-3* and *ros1-7 vs*. Col-0.

Supplemental Table 4: *ROS1*-dependent and *ROS1*-independent maternally-hypomethylated regions.

Supplemental Figure 1: Demethylase expression in seeds and flow cytometry profiles of seed and leaf.

Supplemental Figure 2: Length distributions of differentially methylated regions (DMRs).

Supplemental Figure 3: Average DNA methylation levels of *ros1* endosperm DMRs are similar across biological replicates and alleles.

Supplemental Figure 4: DNA methylation levels at ends of *ROS1* target-associated TEs in endosperm and leaf.

Supplemental Figure 5: DNA methylation levels at ends of TEs unassociated with ROS1 target regions in endosperm and leaf.

Supplemental Figure 6: DNA hypermethylation is limited in ros1 mutant endosperm relative to leaf or sperm.

Supplemental Figure 7: Genome browser displaying 5mC in Col-0, ros1-3, ros1-7 endosperm at the DOGL4 locus (AT4G18650).

Supplemental Figure 8: Hypermethylation in *ros1-1* mutant endosperm is biased for the paternal allele.

Supplemental Figure 9: Hypermethylation in *ros1* mutant endosperm is biased for the paternal allele.

Supplemental Figure 10: DNA methylation levels in additional replicates at maternally-hypomethylated regions.

Supplemental Figure 11: *ROS1*-dependent, biallelically-demethylated regions are hypermethylated in *ros1-3* mutant sperm cells.

Supplemental Figure 12: Fraction of *ROS1* reads that are paternal in origin.

